# Robustness of bristle number to temperature and genetic background varies with bristle type and is regulated by *miR-9a*

**DOI:** 10.1101/295485

**Authors:** A. Matamoro-Vidal, T. Tully, V. Courtier-Orgogozo

## Abstract

Robustness is the invariance of a given phenotype when faced with a given incoming genetic or environmental variation. Such essential property is being studied in a wide diversity of traits in many organisms but it is difficult to compare the results obtained on the robustness of different traits with each other given the differences that exist between traits in the way they are measured, in their genetic architecture and development. In this study, we assessed robustness of bristle number to incoming genetic and environmental variation for eight bristle types in the fruit fly *Drosophila melanogaster,* allowing for a straightforward comparison of the robustness observed between bristle types. We measured the response of bristle number mean and variance to changes in temperature and in the number of copies of two genes (*scute* and *miR-9a*) known to be involved in bristle development. Many combinations between the three factors were tested, thus allowing to test for the effect of each factor in different contexts for the two other factors – to which we refer herein as different backgrounds. We have found different responses between bristle types, suggesting that they present different levels of robustness to the factors tested. In addition, we have found that temperature and *miR-9a* affect more generally the variance of the traits rather than their means, thus fulfilling a criteria usually admitted to identify robustness factors.

## Introduction

Multicellular living organisms are characterized by the development of reproducible visible outcomes from an initial single cell, irrespective of the environmental conditions. A phenotypic trait is considered robust when it displays little or no variation in the face of a particular environmental or genetic perturbation (Debat & David, 2001; Félix & Barkoulas, 2015). The mechanisms that produce robustness are poorly understood. On one hand, specific molecules such as heat shock proteins (Rutherford & Lindquist, 1998), other chaperone proteins (Jarosz & Lindquist, 2010), histones, transcription factors (Gibson & Dworkin, 2004; Siegal & Bergman, 2002) or microRNAs (Vidigal & Ventura, 2015) have been put forward as buffering factors. On the other hand, general features of developmental systems such as redundancies, feedback loops and nonlinearities in quantitative relationships between molecular components, have been proposed to buffer against perturbations and suppress phenotypic variation. Both explanations are not mutually exclusive and can co-exist.

Three types of robustness can be distinguished: robustness to changes in the external environment (such as variation in temperature or in food), robustness to genetic perturbations, and robustness to developmental noise (*i.e.,* noise due to low number of molecules in reacting systems, stochastic gene expression, stochastic cell-by-cell interactions, etc.). Robustness to developmental noise can be assessed in symmetrical traits by measuring fluctuating asymmetry, *i.e.* the deviation from perfect symmetry between left and right sides of the body (Debat & David, 2001). Several experimental studies have attempted to assess whether traits that are robust to one type of perturbation also tend to be robust to other kinds of variations. Overall, mixed results have been obtained and general claims are difficult as it is not possible to test for all possible perturbations. A meta-analysis of knock-out mutations in all non-essential genes in yeasts revealed that genes confering robustness to environmental or stochastic change also buffer the effect of genetic changes (Lehner, 2010). In contrast, the *HTZ1* gene, which encodes a histone variant involved in chromatin regulation, was found to increase robustness to microenvironmental variation but not to random mutations (Richardson et al., 2013). Other studies have compared robustness within species with variation between species. Analysis of cell lineages in various strains of nematodes has thus shown that cell lineages that are more variable within a genetically homogeneous population tend to evolve more rapidly between closely related species than cell lineages that are constant within a strain (Delattre & Félix, 2001).

The bristles covering the body of adult *Drosophila melanogaster* flies form an ideal model system to assess robustness and sensitivity of bristle number to various perturbations. Adult bristles are external mechanosensory organs required for accurate perception, proprioception, locomotion and flight (Kernan, 2007). More than 100 types of bristles can be distinguished on the adult body based on their shape, their number, their stereotyped position and the orientation of their shaft (Lindsley & Zimm, 1992). Each mechanosensory bristle, such as the ones studied in this paper, is composed of four cells including a mechanosensory neuron. The four bristle cells originate during development from a single cell, the sensory organ precursor (SOP) cell, through asymmetric cell divisions (Lai & Orgogozo, 2004). SOP formation is controlled by a network of regulatory factors involving an input/output fate switch mediated by the three transcription factors Achaete (Ac), Scute (Sc), and Senseless (Sens) (Quan & Hassan, 2005; Stern & Orgogozo, 2009). Ac and Sc are bHLH proteins that activate transcription of several target genes including Sens, which feeds back to regulate transcription of the *ac* and *sc* genes (Jafar-Nejad et al., 2003; Nolo et al., 2000). The switch from an epidermal cell to a SOP cell fate involves up-regulation of *ac* and *sc* transcription levels and depends on amplification of subtle initial differences in the transcription of *ac* and *sc* (Quan & Hassan, 2005).

In the 60s, Rendel examined robustness of adult bristle number in *Drosophila melanogaster*. He developed an elaborate model for bristle formation that assumed that the probability of developing a bristle was directly determined by the amount or activity of relevant regulatory molecules, a quantity that he called ‘Make’ (Rendel, 1967). Rendel estimated the hypothetical variable Make from a set of various alleles affecting bristle number and observed little variation from the wild-type phenotype for a range of Make values, and that when Make values go below or over certain thresholds then bristle number either decrease or increase, respectively. Certain bristle types, such as the scutellar and thoracic bristles, were found to be largely invariant or canalized over a large range of Make values, whereas others tended to be more variable.

By comparing how sternopleural bristle number responds to both genetic and environmental perturbations in a panel of naturally derived lines, the mechanisms underlying canalization and developmental stability were inferred to be distinct (Dworkin, 2005). Following Rendel’s seminal study, several authors assessed the possible role of certain candidate genes on robustness of bristle number to various perturbations. Milton et al. (2005) examined 6 bristle categories in *hsp83* mutants and found that *hsp83* alleles and genetic background affect bristle trait means but not variances nor developmental instability between left and right sides of the fly. Transcriptional suppression of h*sp22* and h*sp67Bc* via RNAi was found to affect fluctuating asymmetry in bristle number (Takahashi et al., 2010). On the other hand, *Hsp70Ba* mutants had no effect on fluctuating asymmetry in bristle number but significantly increased among-individual variation (Takahashi et al. 2011). Analysis of a collection of >400 isogenic deficiency lines encompassing approximately 63.6% of the enti*re D. melanogaster* genome identified many deficiencies with significant effects on bristle number mean, on its environmental sensitivity, or on fluctuating asymmetry (Takahashiet al, 2012; Takahashi et al., 2011), suggesting that multiple genomic regions can influence variation in bristle number. Significant correlations for different bristle traits were found among the effects of deficiencies on environmental sensitivity, suggesting that the same genetic mechanisms can regulate environmental sensitivity of various bristles.

An important regulator of robustness of bristle number to genetic and environmental parameter is the gene *mir-9a*. Flies lacking *mir-9a*, which are viable and fertile, sporadically develop extra thoracic (scutellar) and wing margin bristles (Li et al., 2006). miR-9a directly downregulates *Sens* expression via the 3′-UTR of *sens* mRNA. Mutation of miR-9a binding sites in the *sens* mRNA significantly change the variance and the average scutellar bristle number, as well as the sensitivity of the scutellar bristle number to changes in temperature (Cassidy et al., 2013). Importantly, the impact of standing genetic variation on bristle phenotype variation is suppressed by miR-9a regulation of *sens* (Cassidy et al., 2013). Selection for individuals with above-average bristle numbers is more efficient when miR-9a levels are reduced or when the *sens* interaction is impaired. Overall, it suggests that the miR-9a contributes to robustness of the scutellar bristle number against genetic variation.

In this study, we test whether certain types of bristles are more robust than others to perturbations (here temperature changes, genetic mutations or developmental noise), and whether bristles that are robust to one type of perturbation also tend to be robust to other kinds of variations. In addition, we explore the function of *miR-9a* as a factor of robustness: does *miR-9a* act similarly in all bristles or does it have distinct effects on various bristle types? A common approach for investigating the effect of a specific mutation on trait robustness is to introgress the mutation of interest into a reference genetic background through successive crosses (Dworkin, 2005; Milloz, Duveau et al., 2008), so that the comparison of the reference line with the newly-made line gives a direct measure of the effect of the mutation. However, this approach can only reveal the effect of a mutation of interest in a given genetic background and cannot provide a general overview of the effect of the mutation in several genetic backgrounds. In this study, we chose to work with non-isogenic lines and to look for general trends that can be observed with several alleles of the same gene.

## Material and Methods

### Drosophila lines and treatments

The list of the *Drosophila* lines used is given in Table 1. All flies were cultured on standard cornmeal–agar medium in uncrowned conditions at 18° C or 25° C.

**Table 1:**
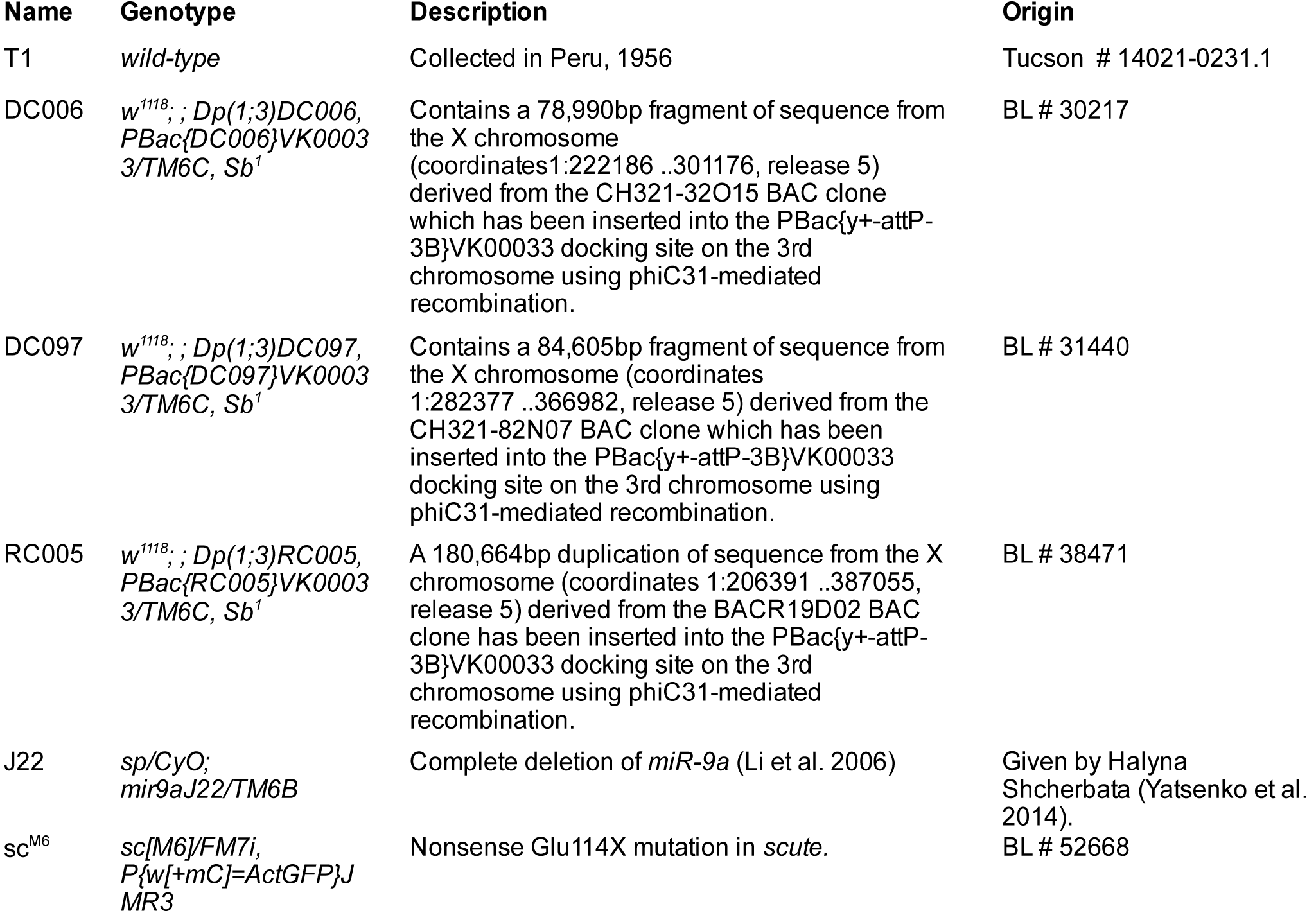
Drosophila lines used. Tucson: San Diego Drosophila Species Stock Center; BL: Bloomington Drosophila Stock Center.

The sample size and genotypes examined for each treatment are shown in Table 2. The gene *scute* is on the X chromosome, so that wild-type males carry a single copy of *scute*. To examine males with two copies of *scute*, males carrying a *scute* duplication (*DC097*, *DC006* or *RC005*) were crossed with T1 virgin females and their male progeny was examined. Males with three copies of *scute* were obtained by selecting within the duplication stock individuals which are homozygous for the duplicated segments. Flies with reduced levels of *scute* were obtained by selecting [non FM7] males from the sc^M6^ stock. To study the effect of removing one copy of *miR-9a* in a variety of *scute* backgrounds, J22 males where crossed with T1, *sc^M6^*, *RC005*, *DC097* and *DC006* virgin females, respectively, and their male progeny was examined. In addition, the effects of depleting two copies of *miR-9a* were tested in three different genetic backgrounds: (1) a background with one copy of *scute* by examining homozygous males from the J22 stock, (2) a background with two copies of *scute* by crossing J22 males with virgin females carrying a recombined 3^rd^ chromosome containing both the *RC005* duplication of the *scute* genomic region and the *miR-9a J22* null allele, and (3) another background with two copies of *scute* by crossing J22 males with virgin females carrying a recombined 3^rd^ chromosome containing both the *DC097* duplication of the *scute* genomic region and the *miR-9a J22* null allele.

**Table 2:**
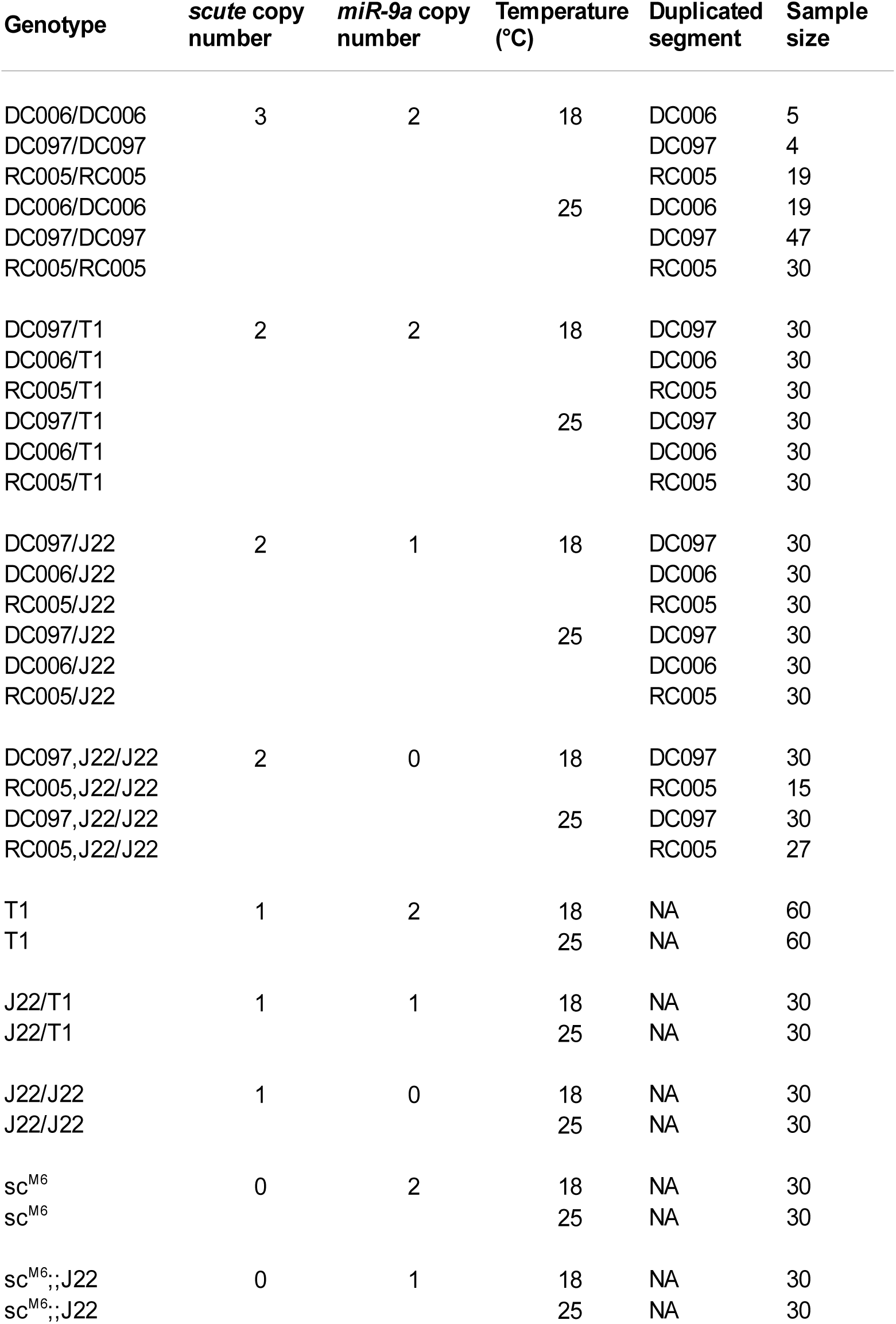
Genotypes investigated and number of flies examined.

**Table 3:** Result of the ANOVA on the *glmer* simplified model for bristle number as a function of bristle type, copies of *miR-9a*, copies of *scute*, temperature (fixed effect terms) *and* scute duplication line (random effect).

|  | Chisq | Df | Pr(>Chisq) |
| --- | --- | --- | --- |
| temperature | 1.09 | 1 | 0.2971 |
| copies.mir9a | 1.42 | 2 | 0.4922 |
| copies.scute | 71.43 | 3 | 0 |
| bristle | 458.98 | 7 | 0 |
| temperature:copies.mir9a | 1.69 | 2 | 0.4285 |
| temperature:copies.scute | 10.89 | 3 | 0.0123 |
| temperature:bristle | 2.57 | 7 | 0.9216 |
| copies.mir9a:copies.scute | 9.12 | 3 | 0.0278 |
| copies.mir9a:bristle | 28.12 | 14 | 0.0137 |
| copies.scute:bristle | 174.58 | 21 | 0 |
| temperature:copies.scute:bristle | 71.54 | 21 | 0 |

Upon collection, flies were stored in 96-well plates in glycerol:acetate:ethanol (1:1:3) solution (one fly per well), allowing long term conservation without risk of breaking bristle shafts.

### Bristles phenotyping

Bristle count was done manually on adult males (n = 946). Bristles were counted by observing the flies in 99% glycerol using a stereomicroscope Carl Zeiss™ STEMI 2000C. All the bristles examined are typically symmetrically located on the left and right sides of the body. We counted bristles for the 8 bristle types illustrated on Figure 1(humeral, dorso-central, mesosternal (Yassin et al., 2007), orbital, vertical, scutellar, ocellar and posterior ocellar).

**Figure 1.**
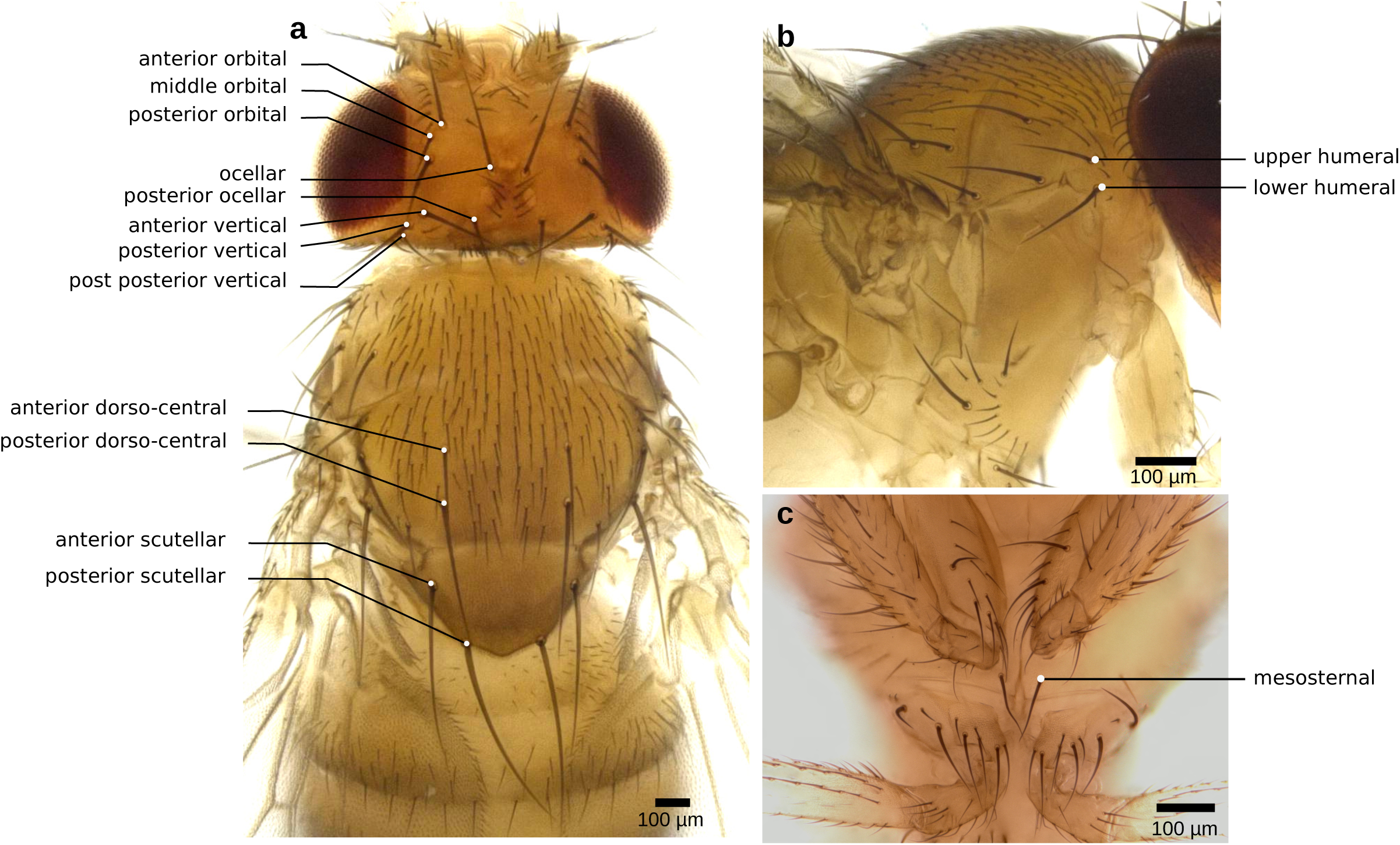
Illustration of the bristles examined in this study. **a:** dorsal view of a *Drosophila melanogaster* T1 adult male raised at 25° C, showing eight head bristles and four thoracic bristles. **b:** lateral view showing the humeral bristles. **c:** ventral view showing the mesosternal bristles. White plain circles indicate the bases of the bristles.

Bristle counts were done by two persons. To verify that both measurements were similar, we compared counts for a set of 36 flies. The set included 12 flies for three genotypes (T1, sc^M6^ and J22/J22), thus totalizing 288 observations (36 individuals x 8 bristles). Congruence between the two counts was then checked. When inconsistencies were found, the corresponding bristles were double checked in order to verify the origin of the discrepancy and make sure that both persons used the same criteria (position, size and orientation of the bristle shaft) to identify each bristle type. 14 observations (4.8%) were incongruous due to confusion between bristle types. The discrepencies concerned the following bristles: vertical (6), orbital (3), mesosternal (1), humeral (3) and dorso-central (1). These inconsistencies disappeared once we designed and used a more precise procedure to identify each bristle type based on position, size and orientation of the birstle shaft.

### Data analyses

#### Bristle number mean and variance.

For each bristle type, bristle number was calculated as the the sum of the bristles located on the left and right parts of the body of each individual. The averagewild-type bristle number is four for scutellar, humeral and dorso-central bristles, two for mesosternal, ocellar and posterior-ocellar, and six for vertical and orbital bristles.

All statistical analyses were performed in R 3.4.1 (R Core Team, 2016). Variation in bristle number was analysed with a Poisson model for count characters. We fitted generalized linear mixed-effect models (GLMM) using the function *glmer* from the R statistical package *lme4* (Wood & Scheipl, 2017), which allows to specify both fixed and random effects. Bristle number was the response variable, and number of copies of miR-9a (0, 1, 2), number of copies of *scute* (0, 1, 2, 3) and temperature (18° C, 25° C) were set as qualitative fixed effect factors. The background for the *scute* duplication lines (*DC006*, *DC097*, *RC005*) was set as a random effect. This choice was guided by the aim of considering general effects of *scute* copy number which are observed over all the backgrounds, and by the fact that in some cases sample sizes are unbalanced between backgrounds (Table 2). We performed model selection, starting by fitting models with all possible interactions between the fixed effects. Non significant effects were removed progressively, starting with the high-level interaction terms, using Type III ANOVA until all the non-significant effects were discarded. We first used a general model on the whole dataset, using bristle type as a covariable. In this model we found significant complex interactions between bristle type and other variables (see results). We thus analysed the effects of *scute*, *miR*-9A and temperature for each bristle type separately.

To better understand the biological meaning of the selected models for each bristle type, we choose to represent graphically the predictions of the selected models. Such choice allows to sense the overall effects of the co-factors and their significant interactions on bristle number but the values are constrained by the selected model. To sense what the data look like without the constraints of the linear model, we used a saturated *glmer* model but without the intercept to derive the predicted means and 95% confidence intervals (CI) for each bristle / treatment combination. Plots of the predicted values for each bristle type are shown on Supplementary Figure 1. We performed pairwise contrasts to test whether changing the temperature, removing one copy of *scute*, or removing one or two copies of *miR-9a* significantly affects bristle number in a variety of genetic and environmental backgrounds. Given the high number of tests done (n = 460), we chose to adjust the p-values using the method of Benjamini & Hochberg (1995), that controls for false discovery rate.

To test whether temperature and the number of copies of *miR-9a* affect the variance of bristle number, we used the function *lme* to fit linear-mixed effect models in which the variance of the residuals can depend on variance covariates using the *varIdent* variance structure options (Pinheiro & Bates, 2000). For each combination of bristle type / temperature / number of copies of *scute*, we fitted a model with the number of copies of *miR-9a* as co-factor and the duplication line background as random effect. Two models were compared: one in which the variance is constrained to be constant, and another one in which the variance was allowed to vary as a function of the number of copies of *miR-9a*. The two models were compared with likelihood ratio tests. In some cases, one of the samples had all the individuals with the same bristle number, which precludes test for variance homogeneity bewteen samples using *varIdent*. For these cases, a test for variance homogeneity was done using Fligner-Killeen’s test, which is a non-parametric test robust to departures from normality. The same procedure was followed to test for an effect of temperature on the variance of bristle number for each combination of bristle type / number of copies of scute / number of copies of *miR-9a*. In total the pairwise constrats allowed to test for the effect of *miR-9a* copy number on bristle number variance in 14 different conditions, and for the effect of temperature on bristle number variance in 9 different conditions for each bristle type.

## Results

In order to assess robustness of bristle number to changes in *scute, miR-9a* and temperature, we decided to study 8 bristle types located on the head and thorax of adult *D. melanogaster* males (Fig. 1). We chose bristles that are symmetrically located on the left and right sides of the body and whose total number is either 2, 4 or 6. We considered as standard conditions the genotype T1 (1 copy of *scute* (X-linked), 2 copies of *miR-9a*) at 25° C. Under these conditions, bristle number mean (+/- standard deviation, n = 60 males) was 4.00 (0) for scutellar and dorso-central bristles; 4.07 (0.25) for humeral bristles; 6.00 (0) for orbital and vertical bristles; 2.0 (0) for ocellar and posterior-ocellar bristles; and 2.12 (0.37) for mesosternal bristles. Thus, all the individuals in our standard conditions have robust phenotypes for all the bristles types (i. e. absence of variance and perfect symmetry), excepted for the humeral bristle where 4 individuals had one additional bristle, and for the mesosternal bristle for which 5 individuals had one additional bristle, and 1 individual had 2 ectopic bristles (one on each side).

### Bristle number is differently affected between bristles types by scute, miR-9a and temperature.

In order to test whether robustness of bristle number to temperature and genetic background varies with bristle type, we counted bristle number for the 8 chosen bristle types in 32 different conditions (Table 2), varying temperature (18°C or 25°C), the number of copies of *scute* (0,1 2 or 3) and the number of copies of *miR-9a* (0, 1, 2). We used a single *glmer* model for all our data, with bristle type, copies of *miR*-9a, copies of *scute*, and temperature as independent variables and we progressively removed the non significant effects, using Type III ANOVA until all the non-significant effects were discarded. The resulting model (Table 3) shows a strong effect of the bristle type term, which is expected because the studied bristle types have distinct average bristle number. We also find a significant interaction between bristle type and copies of *scute* as well as between bristle type and copies of *miR-9a*. Moreover, there is a significant triple interaction between bristle type, temperature and copies of *scute*. These interactions indicate that bristle types differ in their response of bristle number mean to the three factors.

In order to further examine the differences between bristle types in the response of bristle number mean to the three factors examined, we fitted a *glmer* model for each bristle type separately, with copies of *scute*, copies of *miR*-*9a* and temperature as fixed effect terms and *scute* background as a random effect co-factor. For each bristle type, the model predicted bristle number means and standard errors and we performed type III ANOVA on the simplified model to test for the significance of the effect of each co-factor on bristle number mean (Fig. 2).

**Figure 2.**
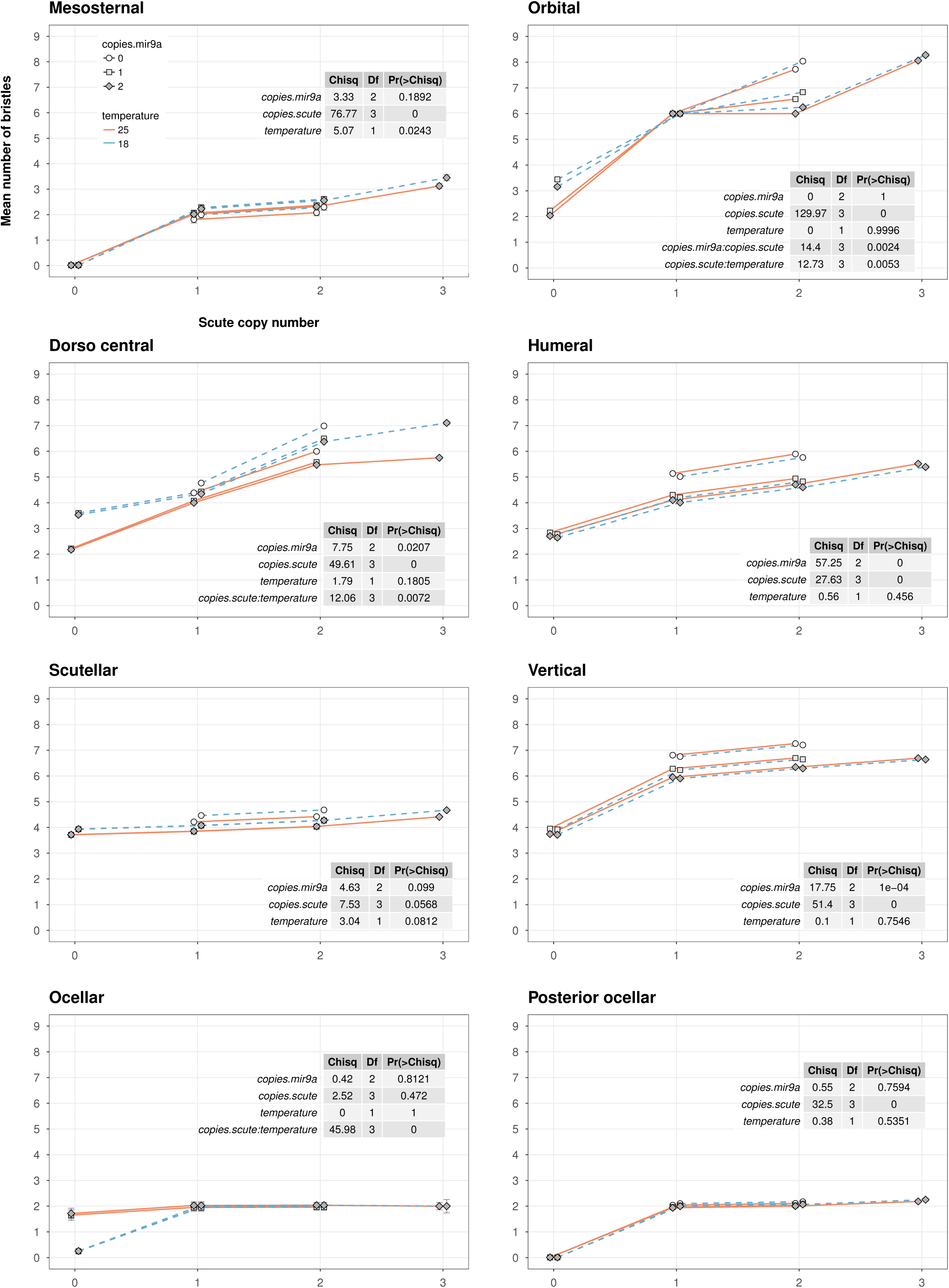
Mean bristle number variation to changes in temperature, *miR-9a* and *scute* copy number. The plots show, for each bristle type, the estimates of bristle number means and standard errors obtained by fitting a simplified generalized linear mixed model for bristle number as a function of temperature, *scute* and *miR-9a* copy number. The results for each bristle type are shown on a separate plot. Within each plot, the results of the type III ANOVA test on the coefficients of the simplified model are shown.

The temperature term was significant by itself or in interaction with another term for 4 out of 8 bristle types. There was no temperature effect for the scutellar, humeral, posterior-ocellar and vertical bristles. For the mesosternal bristle, temperature descreases bristle number in all the conditions excepted when copies of *scute* is 0. For the dorso-central bristle, temperature generally decreases bristle number, with less pronounced effects when copies of *scute* is 1. For the orbital bristle, temperature also decreases bristle number but only when copies of *scute* is 0. Notably, the ocellar bristle at copies of *scute* 0 is the only case where temperature increases bristle number, with more bristles at 25 °C than at 18 °C. Overall, the effect of temperature greatly varies depending on bristle type, and *scute* copy number, but does not interact with *miR*-*9a* copy number.

The effects of *scute* was significant for 7 bristle types. The scutellar bristle was the only bristle not affected by this factor, although the term is marginally significant (p ~ 0.057). In all cases, as expected, the number of bristles increases with the number of copies of *scute.* Three bristles (vertical, ocellar and posterior-ocellar) are affected only by depleting *scute*, and this effect was totally dependent on the temperature for the ocellar bristle only. For mesosternal and orbital bristles, bristle number is mostly affected by extreme values of copies of *scute* (from 0 to 1, and from 2 to 3 copies), although for the orbital bristle the effect of *scute* is dependent on the number of copies of *miR*-*9a*. Finally, the dorso-central and the humeral bristle number increase roughly linearly with the *scute* copy number. Although *scute* copy number affects almost all bristles, the response observed for each bristle is very different with respect to the magnitude of the effects and the interactions with the two other co-factors (temperature and *miR-9a)*.

No effect of *miR-9a* copy number was detected for the scutellar, mesosternal, ocellar and posterior-ocellar bristles. For the other bristles, decreasing the number of copies of *miR*-*9a* increases bristle number. For orbital and vertical bristles, depleting one copy of the gene was sufficient to observe an effect, and the effect was stronger by totally depleting the gene. The effect of *miR-9a* was strongly dependent on *scute* copy number for the orbital bristle type. Finally, the dorso-central and the humeral bristle number were affected only by total depletion of the *miR-9a* gene. As in the case of temperature and *scute,* the effect of *miR-9a* thus differs between bristle types.

Figure 3 summarises for each bristle type the factors or combination of factors that were found to affect bristle number mean. The response of each bristle type is unique except for the vertical and humeral bristles which are sensitive to the same factors.

**Figure 3.**
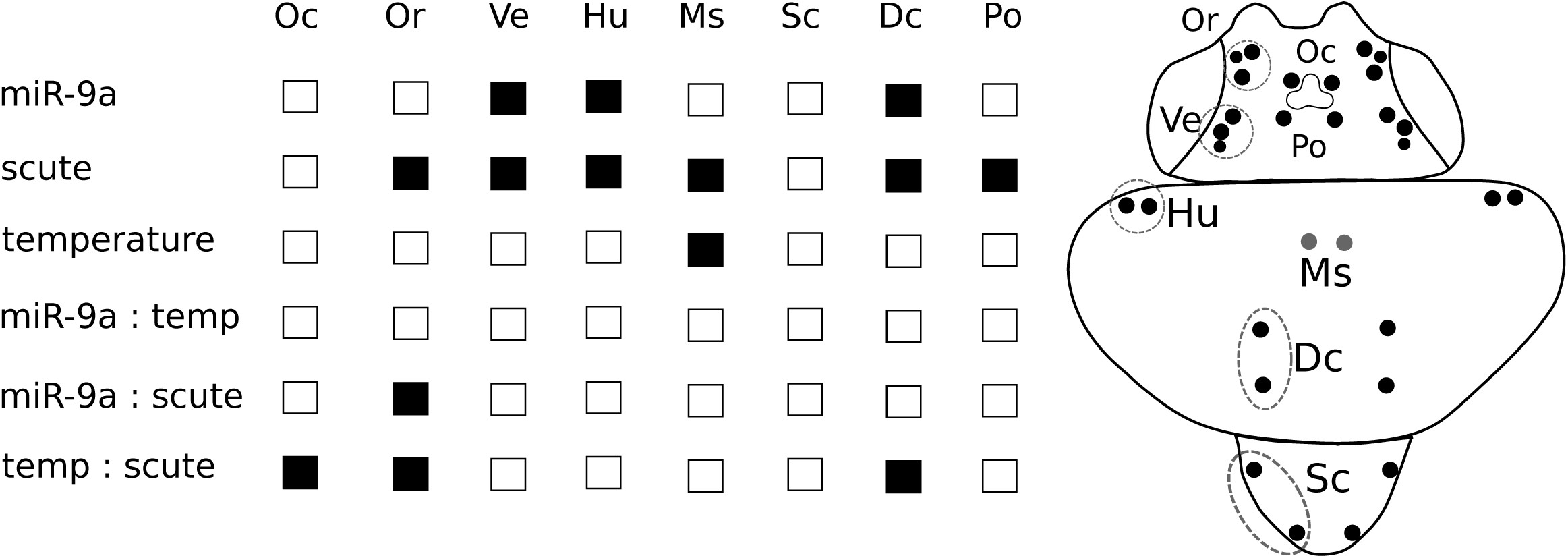
Summary of the type III ANOVA tests performed on the simplified generalized mixed models. All the studied factors and their interactions are shown. For each bristle type, black filled squares indicate the factors or interaction of factors having a significant effect on bristle number mean. White squares indicate factors or interaction of factors that did not have a significant effect on the bristle number mean. A diagram of a fly dorsal view shows the location of each bristle type. Oc, ocellar; Or, orbital; Ve, vertical; Hu, humeral; Ms, mesosternal; Sc, scutellar; Dc, dorso-central; Po, posterior ocellar.

To assess the effects of each factor in a variety of conditions for the two other factors, we performed pairwise comparisons and evaluated the impact on bristle number of varying one co-factor having all the other co-factors fixed. We thus tested the effects of varying *scute* copy number (from 0 to 1; 1 to 2; or 2 to 3 copies), *miR*-*9a* copy number (from 0 to 1, from 1 to 2 and from 0 to 2), and temperature (from 18 °C to 25 °C) all things equal otherwise.

The effects on bristle mean of changing one copy of *scute* were studied in 12 different combinations of *scute* copy number, *miR-9a* copy number and temperature in 8 bristle types (supplementary table 1). On 96 tests performed, 37 were significant and pertained to all the bristle types excepted the scutellar bristle (Fig. 4). The effects of *scute* copy number were observed in a variety of backgrounds and no specific patterns could be detected (supplementary table).

**Figure 4.**
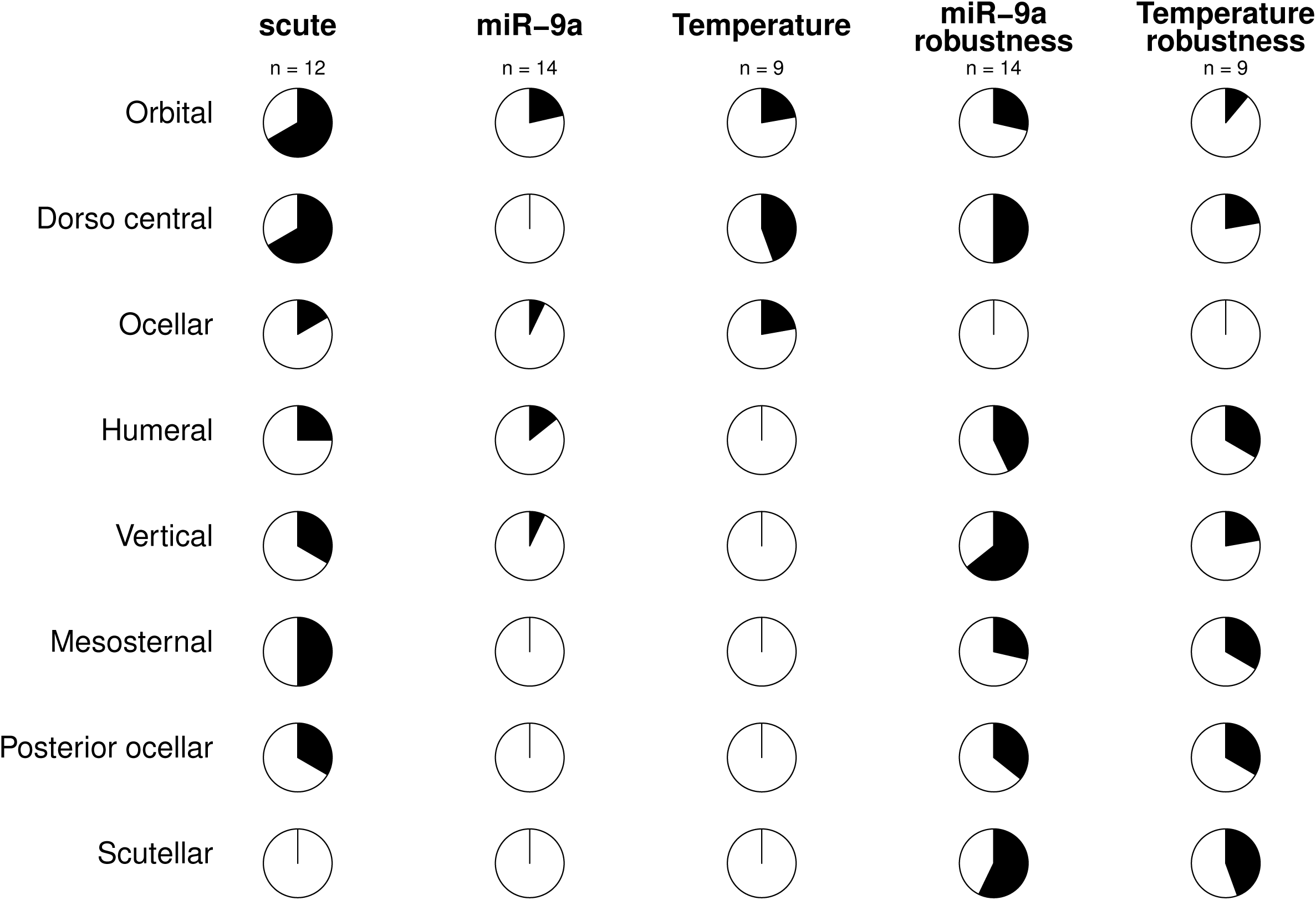
Summary of the results for the pairwise contrasts. The first three columns of pie charts show the proportion of conditions for which there was a significant effect on bristle number mean further to changes in *scute* copy number, *miR-9a* copy number, and temperature respectively. The fourth and fifth column of pie charts show the proportion conditions tested for which there was significant effect on bristle number variance but not on bristle number mean, further to changes in *miR-9a* copy number, and in temperature respectively.“n” is the number of conditions tested.

For *miR-9a*, the effects on bristle mean of changing one or two copies of *miR-9a* were studied in 14 backgrounds for each bristle type thus totalling 112 tests. We found effects on bristle mean in 7 tests only, representing 4 bristles (Fig. 4, supplementary table 1). An effect on bristle mean of *miR-9a* copy number was observed in 3 different backgrounds for the orbital bristle, in 2 backgrounds in the humeral bristle and in one background only for the ocellar and vertical bristles. In all cases except one (the effect of *miR-9a* on humeral bristles at 25 ° C), the effect of *miR-9a* on bristle mean was observed in a non wild-type background and/or at 18 ° C.

For the temperature, we found a significant effect on bristle number mean for 8 out of 72 conditions tested (9 conditions tested for each bristle). The effects were observed mostly in the dorso-central bristle (4 cases), and in the orbital and ocellar bristles (2 cases each) (Fig. 4). An effect of the temperature on bristle number mean was observed only in non wild-type conditions (i.e., when *scute* copy number is not 1 and/or *miR-9a* copy number is not 2).

This second analysis confirms that bristle types differ in their sensibility to the three various factors, with certain bristles being more sensitive to certain factors than others. Furthermore, we found no correlation between the effects of the three factors: increased sensibility to one parameter does not appear to be associated with increased sensibility for another factor. Finally, this analysis shows that *scute* affects bristle number in almost all the conditions considered (although in a different fashion depending on the bristle type), whereas the effects of *miR-9a* and temperature are more scarce, and observed preferentially in conjonction with changes in other factors.

### miR-9a and temperature affect bristle number robustness.

It has been suggested that *miR*-*9a* is a robustness factor protecting phenotypic variation from incoming genetic or environmental perturbations (Cassidy et al., 2013). Strictly speaking, a robustness factor modifies the variance of the phenotype without changing its mean (Félix & Barkoulas, 2015). To test whether *miR-9a* acts as a robustness factor for bristle number in the 8 studied bristle types, we used our pairwise contrast tests on bristle means and variance to search for cases where *miR-9a* copy number affects bristle number variance without changing bristle number mean. We measured the effects on mean and variance of varying *miR-9a* copy number in 14 different genetic and environmental contexts (Supplementary Table 1). We found a significant effect of *miR-9a* copy number on bristle number variance but not on bristle number mean in 48 out of 112 tests (14 backgrounds x 8 bristles). We observed changes in bristle number variance without changes in the mean in at least 4 out of 14 backgrounds for all the bristles excepted for the ocellar bristle which was not affected at all (Fig. 4). Interestingly, for 5 bristles (humeral, mesosternal, posterior-ocellar and scutellar), an effect of *miR-9a* copy number on bristle number variance without any effect on bristle number mean was never seen in the wild-type conditions (1 copy of scute, 25° C). For these bristles, the effects of *miR-9a* on the variance occur only in conjunction with a change in another factor (temperature or *scute* copy number). For the dorso central bristle, *miR-9a* copy number variation affected bristle number variance but not the mean in 7 different backgrounds, two of which were wild-type (Supplementary Table 1). For the vertical bristles, *miR-9a* copy number affected bristle number robustness in 9 conditions, but only two of them were wild-type (Supplementary Table 1). Overall, these data show that *miR-9a* is more likely to affect bristle number variance than bristle number mean, and that these effects are observed preferentially in conjunction with changes in other factors.

We also measured the effect of temperature on bristle number robustness by searching conditions where temperature affects bristle number variance without affecting the mean. This was done in 9 different genetic backgrounds for each bristle type thus totalling 72 tests. We found a significant effect of the temperature for the bristle number variance and not for the mean in 18 tests, corresponding to all the bristles excepted the ocellar bristle (Fig. 4). Strikingly, out of the 18 significant effects of the temperature observed, only one was detected in wild-type conditions and all the other ones were seen in conjunction with variation in other factors.

Our data thus show that *miR-9a* copy number and temperature rarely impact the bristle number mean and they tend to affect more broadly bristle number variances. In addition, we found that the effects on bristle number variance are rarely observed in standard wild-type conditions and require a sensitive background.

## Discussion

Understanding the response of phenotypic variation to mutations and to environmental perturbations is important to explain phenotypic evolution. For example, the pattern of mutational phenotypic variation observed at the microevolutionary level (i.e., within species) can be a good predictor of the pattern of variation observed at the macroevolutionary level (between species) (Farhadifar et al., 2016; Houle et al., 2017),. In this study, we examined whether we could find differences in the robustness of bristle number to genetic and environmental perturbations between 8 bristle types in *D. melanogaster* which, in principle, share a similar genetic architecture and underlying developmental gene network.

Differences in robustness to genetic and environmental incoming variation were assessed by measuring the response of bristle number mean and variance to three parameters: the number of copies of the gene *scute*, the number of copies of the gene *miR-9a,* and temperature, with many combinations between the three factors. This allowed us to study the effect of one of the factors in several different contexts. To increase the number of copies of *scute*, we used lines carrying duplicated segments of the X chromosome encompassing both the coding region of *scute* and flanking regions containing many known regulatory regions involved in the patterning of the bristles under study (Campuzano et al., 1985). However, it is possible that the duplications studied do not contain all the cis-regulatory regions for the 8 bristles. If cis-regulatory regions are missing for some bristle types, then varying the number of copies of *scute* will affect the amount of *scute* differently for these bristles, thus creating differences between the bristle types in their response to an increase of *scute* copy number. To study the effect of decreasing *scute* levels, we used a null nonsense coding allele of the gene, which in principle should affect the expression of *scute* in a similar fashion for the 8 bristle types.

In general, the number of bristles tended to increase with the number of copies of *scute,* although the relationships between the two variables differed between the bristle types. As explained above, differences between bristles in their response to increased levels of *scute* could be due to the possibility that for some bristles the duplication lines used didn’t encompass their corresponding regulatory regions. We also observed differences between the bristle types in their response to decreased levels of *scute*. For the scutellar bristle, the ocellar bristle at 25° C, and the dorso-central bristle at 18° C, bristle number was not significantly reduced in the *sc[M6]* genotype although it was for all the other bristle types. These data suggest that some bristles are less sensitive than others to *scute* decreased levels. In all our tests, we found no effect of *scute* on scutellar bristles (Fig. 2-4). Since the cis-regulatory regions responsible for scutellar bristle development are included within the *DC097* and *RC005* duplications we used (Campuzano et al., 1985), the insensitivity of the scutellar bristles to levels of *scute*, compared to the other tested bristles, must be due to differences in their gene network underlying bristle number robustness.

We studied the effect of *miR-9a* copy number using a loss-of-function allele in which the *miR-9a* precursor DNA sequence is entirely deleted (Li et al., 2006). Such allele is expected to have a similar effect on *miR-9* expression levels in all the bristle types. Three bristles only (dorso central, humeral and vertical) were affected by *miR-9a* by itself (Figs. 2-3). In all cases, bristle number increased with reduced levels of *miR-9a*, consistent with the known effect of *miR-9a* as a repressor of Senseless expression (Li et al., 2006), a protein necessary for the differentiation of sensory organ precursor cells. Thus, the studied bristles are not equally sensitive to *miR-9a* copy number.

Regarding temperature, we found that it affects bristle number mean for the mesosternal bristle only. In addition, temperature affects bristle number mean in interaction with *scute* copy number for three other bristles (orbital, ocellar and dorso central). In general, we observed that an increase of the temperature decreases the number of bristles, with the exception of the ocellar bristle at 0 copies of *scute*, where the number of bristles increases with the temperature. As in the case of *scute* and *miR-9a*, the effect of the temperature greatly varies with the bristle type.

One way to identify a robustness factor consists in monitoring whether variation in this factor affects the variance of the trait without affecting its mean (Félix & Barkoulas, 2015). We found here that both the number of copies of *miR-9a* and the temperature affect bristle number variance but not the mean. We monitored for each bristle type whether changes in *miR-9a* copy number and in temperature affected bristle means in a variety of backgrounds. We found that the means of bristle number were affected in relatively few cases (Fig. 4). In most of them, the effects were observed in non wild-type conditions (supplementary table). With the exception of the ocellar bristle type, all the bristles responded to variation in *miR-9a* copy number and in temperature by changes in their bristle number variance without changes in bristle number mean (Fig. 4). Interestingly, changes in bristle number variance were observed mainly in non wild-type conditions. Thus, although *miR-9a* and temperature by themselves have only slight effects on bristle number means, they do have general effects on bristle number variances. Particularly striking is the case of the scutellar birstle, whose bristle number mean was found to be totally robust to changes in *scute,* temperature and *miR-9a* levels (Figs. 2-3) but which bristle number variance was affected by temperature and *miR-9a* copy number in many genetic and environmental contexts, confirming the role of *miR-9a* as a robustness factor in the context of the scutellar bristle development (Cassidy et al., 2013; Cohen et al., 2006).

In conclusion, our data suggest that despite being similar organs, the different bristle types have evolved different gene networks such that all the bristles do not respond in the same way to genetic and environmental perturbations. Such differences could lead to variation in the evolutionary capacity between bristle types. In addition, we found that *miR-9a* acts as a general robustness factors on all the bristle types we examined, thus generalising previous findings on individual bristles.

## Acknowledgments

We thank Loïc Bodet and Mathilde Notin for gathering the dataset; Halyna Shcherbata, the Drosophila Bloomington Stock Center and the San Diego Drosophila Species Stock Center for sending us flies, and Marie-Anne Félix and all the members of the Courtier-Orgogozo lab for stimulating discussions.

## Supplementary Material

**Figure S1:**
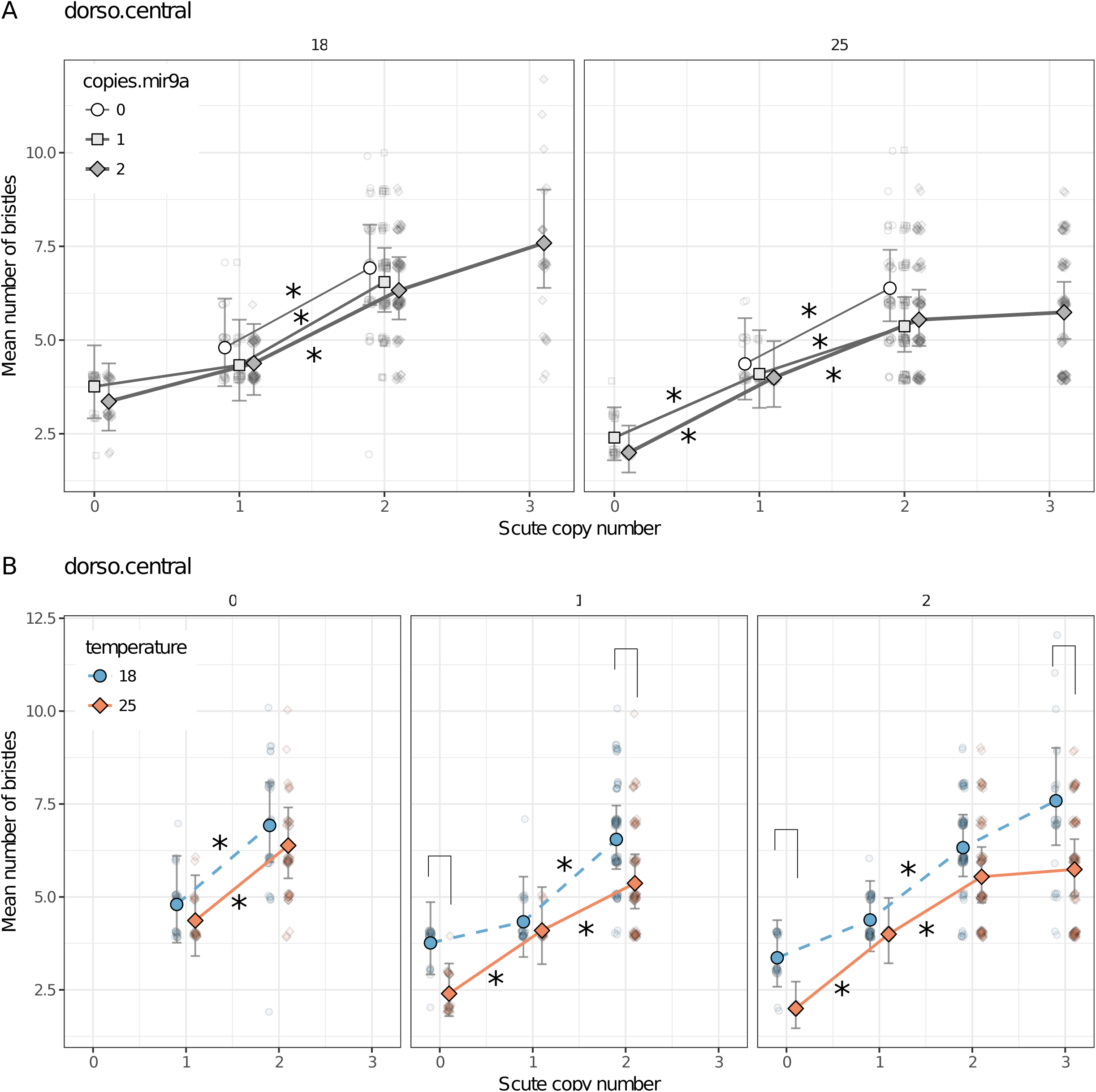
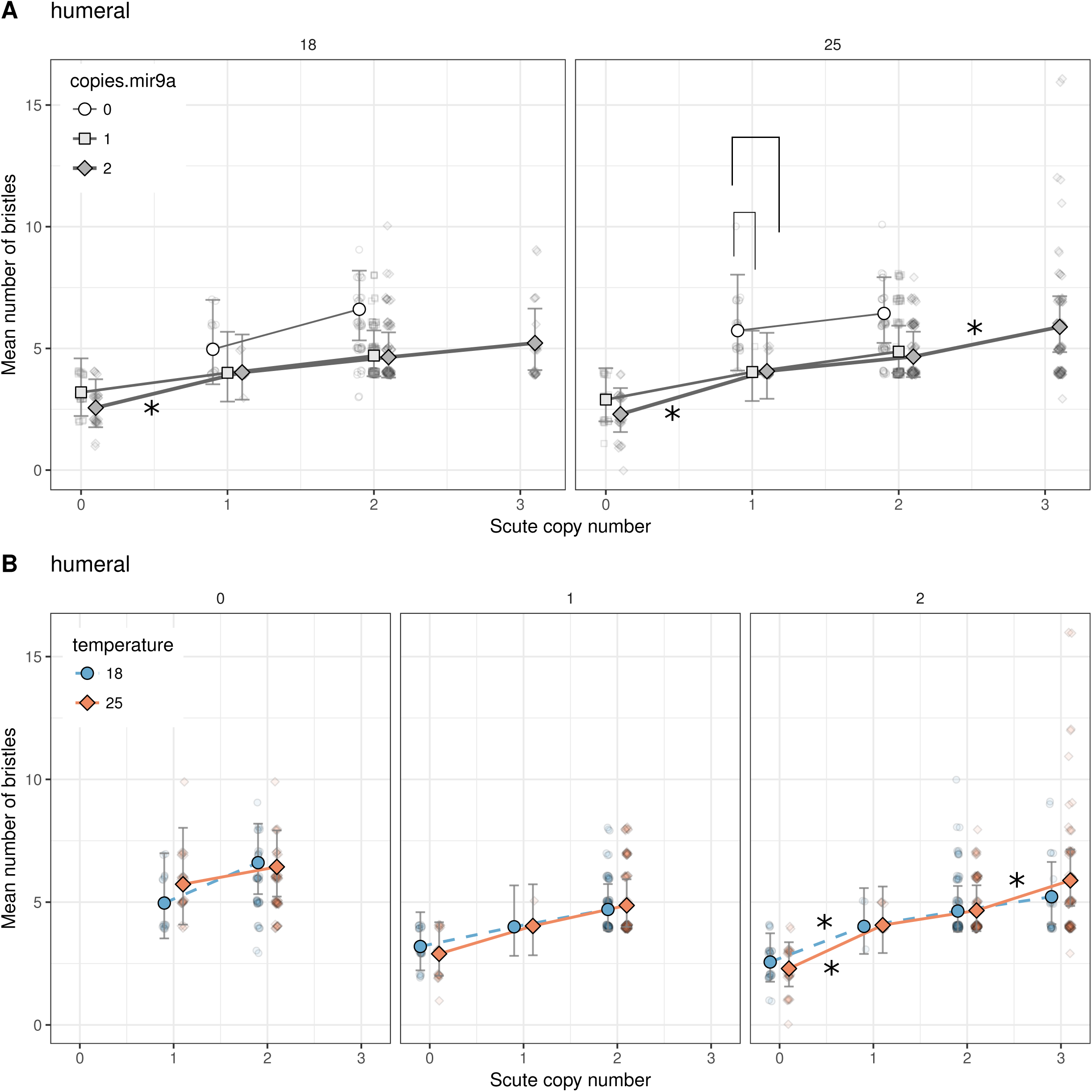
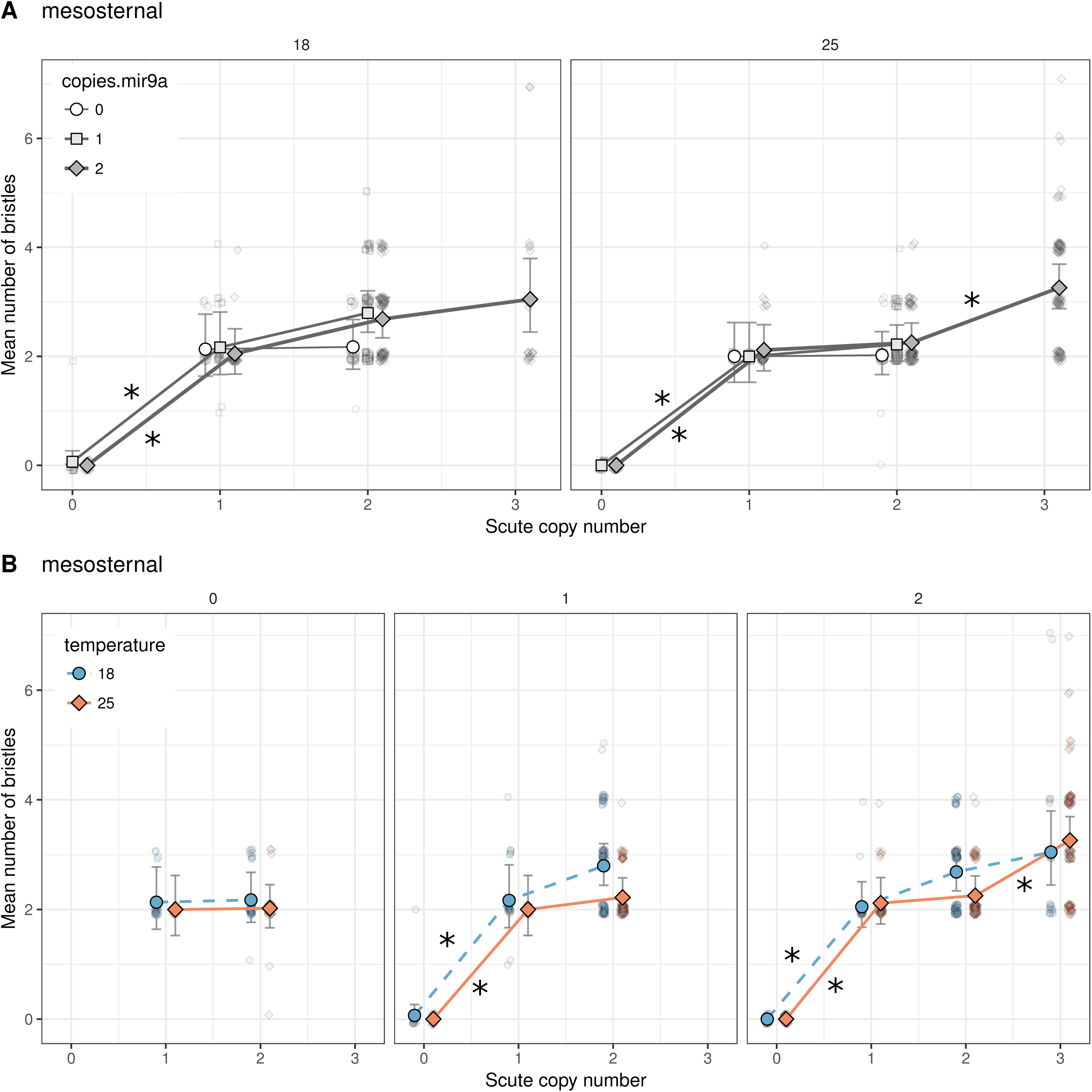
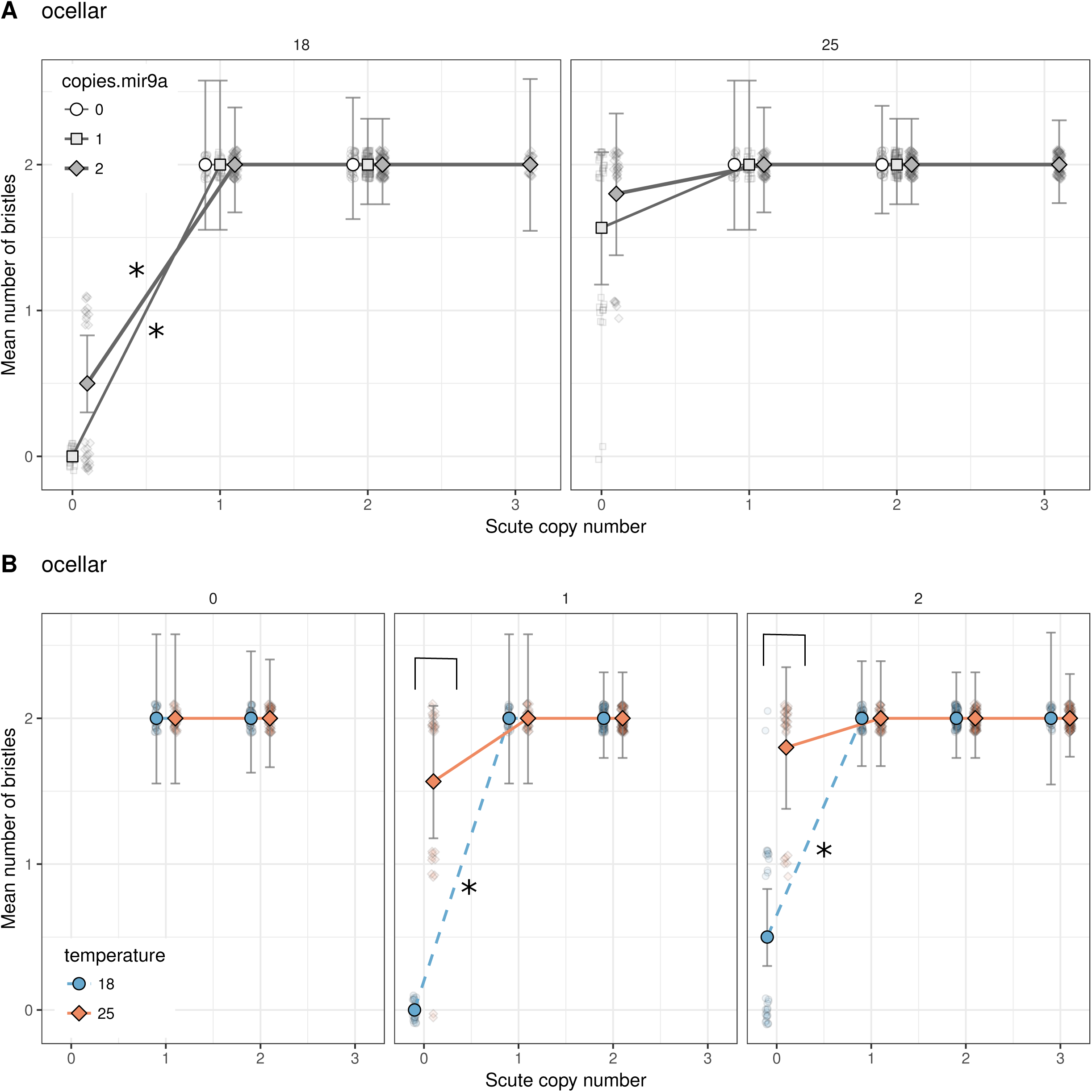
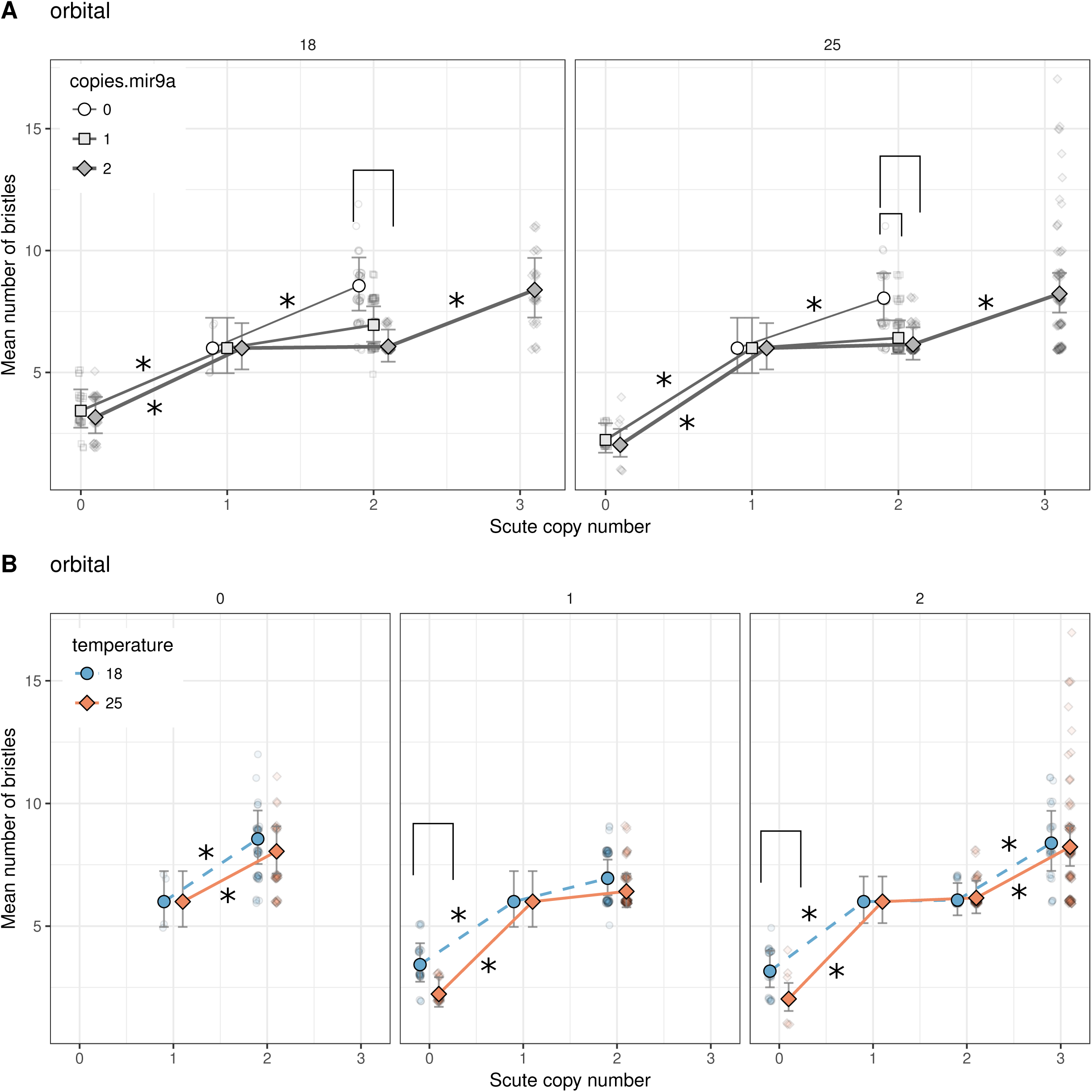
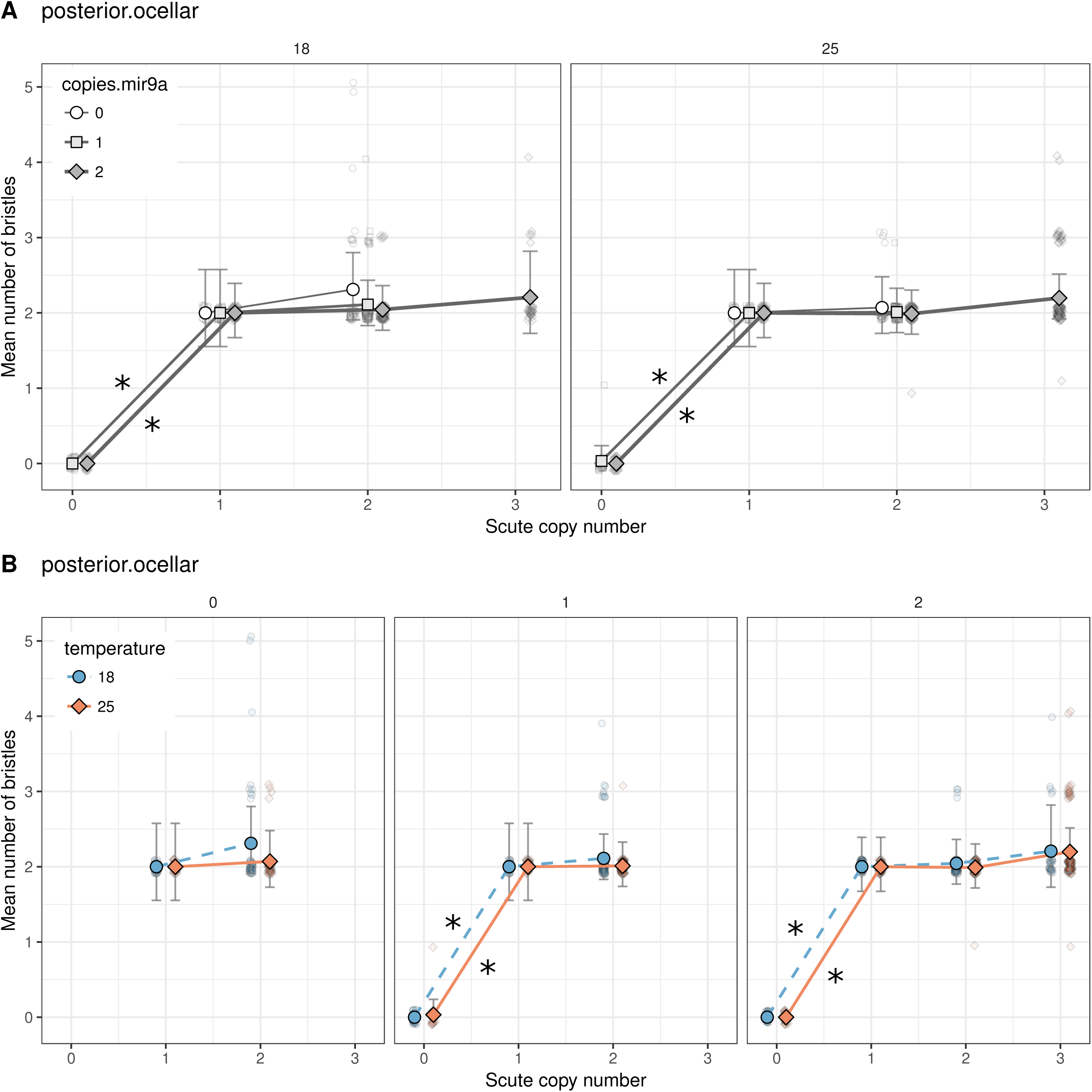
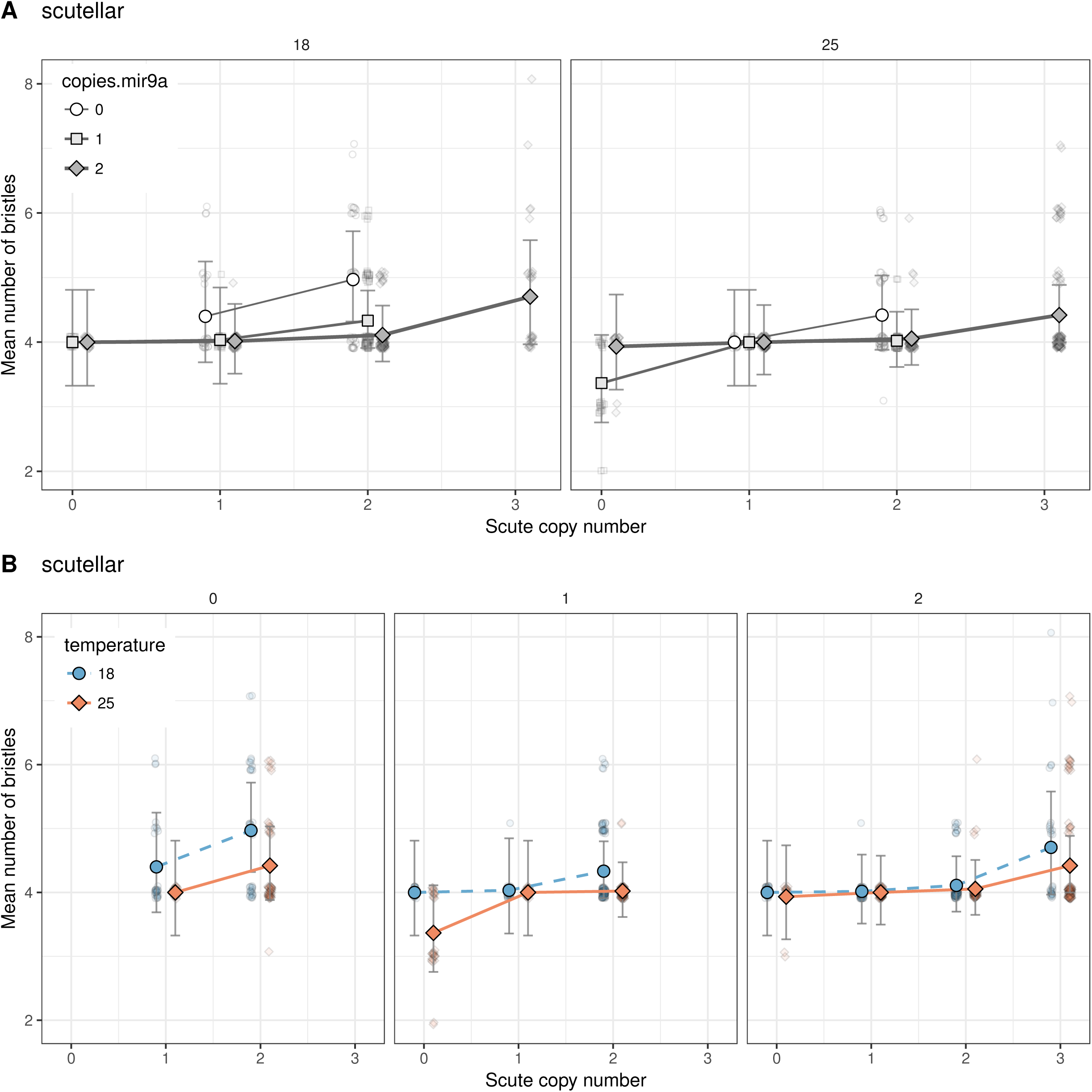
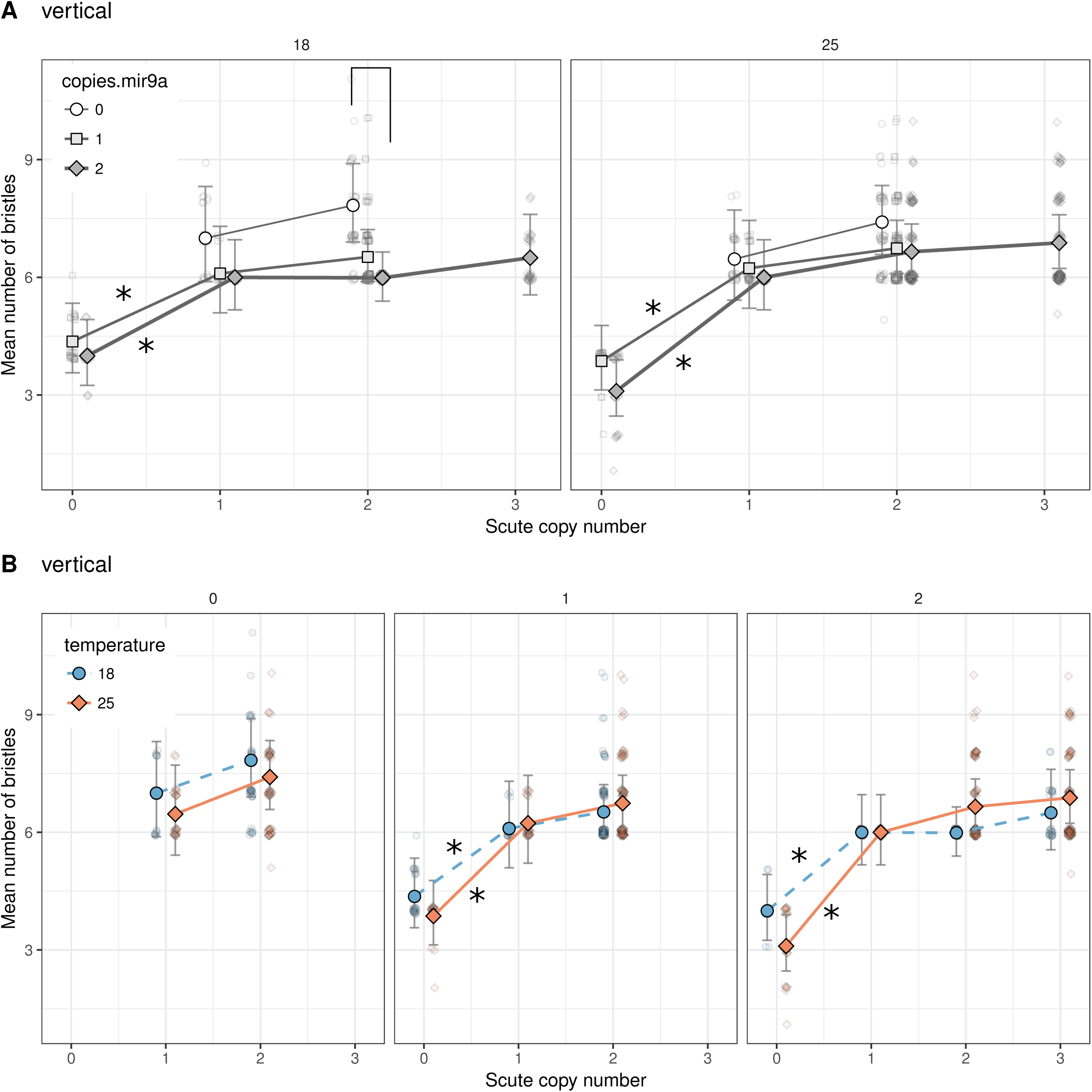
Predicted values for bristle means and 95% confidence intervals using a saturated *glmer* model but without the intercept. The response of bristle number means for each bristle type is shown with emphasis on the variation of *miR-9a* copy number (top pannels) and with emphasis on the variation of temperature (bottom pannels). “*” on lines indicate pairwise comparison for which there was a significant effect of *scute* copy number on bristle number (p < 0.05). Brackets indicate pairwise comparisons for which there was a significant effect of *miR-9a* copy number or of temperature on bristle number (p < 0.05). Light symbols show the raw data values.

**Supplementary table 1. Results for the pairwise contrast tests.** Each row represents a combination of values for the variables bristle type; temperature; number of copies of *miR-9a* and number of copies of scute. **P scute** is the p-value of the effect on bristle number mean of removing one copy of *scute* relatively to the number of copies of *scute* shown on the corresponding row. For example, the value of P scute at a row where copies of *scute* is equal 2 indicates the significativity of the test comparing bristle number between the condition where copies of *scute* is equal 2 and the condition where copies of *scute* equals 1 all things equal otherwise. **T temp** is the p-value linked to the effect on bristle number mean of changing the temperature from 25° C to 18° C all things equal otherwise. **P miR-9a** is the p-value of the effect on bristle number mean of varying one or two copies of *miR-9a* relatively to the number of copies of miR-9a shown on the corresponding row. At rows where copies of *miR-9a* is equal 2, P miR-9a indicates the significativity of the test comparing bristle number mean between the condition where copies *miR-9a* is 2 and the condition where copies of *miR-9a* is 1. At rows where copies of *miR-9a* is equal 1, P *miR-9a* indicates the significativity of the test comparing bristle number mean between the condition where copies of *miR-9a* is 1 and the condition where copies *miR-9a* is 0. At rows where copies miR-9a is equal 0, P miR-9a indicates the significativity of the test comparing bristle number mean between the condition where copies of *miR-9a* is 0 and the condition where copies of *miR-9a* is 2. **P var miR-9a** is the is the p-value of the effect on bristle number variance of varying one or two copies of miR-9a relatively to the number of copies of miR-9a shown on the corresponding row. This row reads in the same way than P miR-9a. **P var temp** is the is the p-value of the effect on bristle number variance of of changing the temperature from 25° C to 18° C. **Robustness miR-9a** denotes the cases for which varying one or two copies of *miR-9a* has a significant effect on bristle number variance (p < 0.05) but not on bristle number mean. **Robustness temperature** denotes the cases for which varying the temperature has a significant effect on bristle number variance (p < 0.05) but not on bristle number mean. NAs denote the redundant cases.

